# Synthetic augmentation of cancer cell line multi-omic datasets using unsupervised deep learning

**DOI:** 10.1101/2024.06.26.600742

**Authors:** Zhaoxiang Cai, Sofia Apolinário, Ana R. Baião, Clare Pacini, Miguel D. Sousa, Susana Vinga, Roger R Reddel, Phillip J. Robinson, Mathew J. Garnett, Qing Zhong, Emanuel Gonçalves

## Abstract

Multi-omic characterization and integration remains a challenge due to data complexity and sparsity. Addressing this, our study introduces an unsupervised deep learning model, MOVE (Multi-Omic Variational Encoder), specifically designed to integrate and augment the Cancer Dependency Map (DepMap). Harnessing orthogonal multi-omic information, this model successfully generates molecular and phenotypic profiles, resulting in an increase of 32.7% in the number of multi-omic profiles and thereby generating a complete DepMap for 1,523 cancer cell lines. The synthetically enhanced data increases statistical power, uncovering less studied mechanisms associated with drug resistance, and refines the identification of genetic associations and clustering of cancer cell lines. By applying SHAP for model interpretation, MOVE reveals multi-omic features essential for cell clustering and biomarker identification related to drug and gene dependencies. This understanding is crucial for the development of much-needed, effective strategies in prioritizing cancer targets.

## Introduction

The growing molecular and phenotypic characterization of cancer cell lines makes them one of the most studied human cell models ^1^. This ever-growing and rich multi-omic data continues to drive the identification of cancer genes and the discovery of novel therapeutic targets ^2–4^. While genomics have dominated the search for predictive biomarkers in cancer, as part of the Cancer Dependency Map consortium recent functional genetic screens showed that mutations and copy number alterations explained less than 20% of RNAi cancer dependencies that could be predicted with molecular markers ^5^. This raises the importance of developing holistic machine learning models capable of integrating, beyond genomics, other types of omics.

Multi-omic integration faces several limitations, most importantly, high heterogeneity of different data types (e.g., discrete vs. continuous distributions), intrinsic technological limitations (e.g., missing values), and limited data availability (e.g., in this study, only 25.8% of the cancer cell lines have a complete set of all seven omic datasets under consideration) ^6^. Unsupervised machine learning has been successful in multi-omic integration, capturing patterns of data variation shared across different omics ^7,8^. This approach highlighted cancer cellular states associated with epithelial-to-mesenchymal transition (EMT), a key process in drug resistance and metastasis ^9^. Unsupervised deep learning-based models can generate improved versions of input datasets, by reconstructing missing measurements and correcting experimental error, and thereby augment downstream analysis ^10,11^. Although linear dimensionality reduction models ^8,12^ have been designed for similar purposes, the application of deep generative models to large-scale multi-omic cancer cell models is lagging behind. This leaves a gap in the utilization of these non-linear approaches to augment the datasets and perform statistical analysis to improve the characterization of cancer mechanisms, novel drug targets and biomarkers ^5,13,14^. Deep learning models, such as variational autoencoders (VAE), provide more complex formulations of the underlying biological data. Moreover, VAEs have highly flexible designs that can handle data sparsity robustly and are easily extensible to incorporate different data types.

Here, we developed a Multi-Omics Variational Auto-Encoder (MOVE) that synthetically augments multi-omic datasets across >1,500 cancer cell lines part of the Cancer Dependency Map (DepMap). MOVE provides a generative unsupervised deep learning model for cancer discovery, using SHapley Additive exPlanations (SHAP) values ^15^ for model explainability that can facilitate the discovery of new biological mechanisms and drug targets. We systematically evaluated and benchmarked MOVE, demonstrating its capabilities in reconstructing independent drug response and proteomic datasets, recovering cancer cell line tissue and cell type clustering, and increasing statistical power to find genomic associations with CRISPR-Cas9 gene essentiality screens. With MOVE, we generated a complete multi-omic profile across all seven different omics, increasing in 32.7% the number of available samples.

## Results

### Unifying deep generative model for cancer multi-omics

Taking advantage of the Cancer Dependency Map (DepMap) project ^5,16,17^, we assembled seven different cancer cell line datasets, i.e., genomics ^2,3^, methylomics ^18^, transcriptomics ^19^, proteomics ^9^, metabolomics ^20^, drug response ^2,18,21,22^, and CRISPR-Cas9 gene essentiality ^4,23^ ***(Figure 1a)***. This comprises a total of 1,523 cancer cell lines for which at least two datasets were available ***(Supplementary Table 1)***. We designed MOVE tailored to the cancer cell lines multi-omic datasets, performed robust data augmentation. and provided model explanations for biomarker discovery (***Figure 1b***, see ***Methods***).

**Figure 1.**
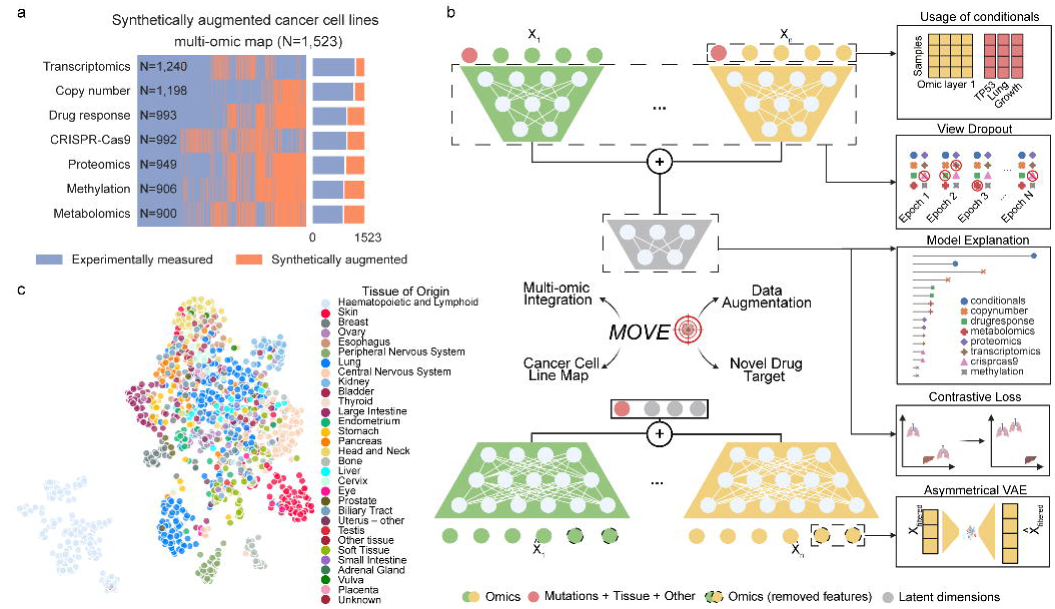
Cancer multi-omic integration with MOVE. **a)** cancer cell line multi-omic datasets integrated. Purple represents measured datasets, while orange represents gaps, i.e., missing datasets, across the 1,523 cancer cell lines. **b)** schematic of the autoencoder, MOVE, where encoders are represented at the top and decoders at the bottom. For simplicity, the integration of only two datasets is represented. Highlighted designs of MOVE are illustrated on the right. **c)** dimensionality reduction visualized using Uniform Manifold Approximation and Projection (UMAP) representation of the trained MOVE joint latent space, where each dot represents a cancer cell line colored according to its tissue of origin.

First, following a late integration ^24^ approach, we trained a separate encoder for each dataset to derive latent embeddings specific to each omic. These embeddings were then concatenated to formulate a joint multi-omic latent representation ***(Figure 1c, Supplementary Table 2)***. Here, a latent representation is a learned, abstracted feature set (embeddings) within the hidden layers of a deep neural network that encapsulates the major information from the input data. Compared to a multi-omic linear dimensionality reduction method, MOFA ^8,12^, our model provides better separation of cell lines by tissue (***Supplementary Figure 1a, 1b***).

Second, genomics presents a unique challenge due to the sparsity and qualitative nature of its data. To address this, we use only frequent cancer driver events and split genomics into copy number events and mutations. While copy number events are integrated as ordinal data through a separate encoder/decoder akin to other omics, mutations are integrated as conditionals to each encoder (***Figure 1b,*** *see **Methods***). The rationale is that the genetic backgrounds influence cellular profiles and phenotypes, thereby conditioning other omic layers. The conditional matrix contains genetic alterations in cancer driver genes (including gene fusions), cell line tissue of origin, cell line growth rate measurements, and microsatellite instability (MSI high), totaling 237 binary conditional variables ***(Supplementary Table 3)***. This conditional matrix is further concatenated to the trained joint latent space that works as input for the decoders. Hence, in MOVE, cell genetic background and phenotypic information play a crucial role in the generation of latent representations and the reconstruction of each omic dataset.

Third, compared with similar models for single cell data ^10,25,26^, the limited number of samples and heterogeneity among all omics pose a challenge to train a generalizable model for cancer cell lines. To reduce model complexity, MOVE only considers the most variable features as input for the encoders, while all features are reconstructed by the decoders for synthetic data generation, resulting in an asymmetrical design of VAE ***(Figure 1b, Supplementary Figure 2a, 2b, Supplementary Table 4)***. This unique design of MOVE allows us to discard low informative features, such as genes with constant expression and non-essential genes across all cancer cell lines. This reduces the number of trainable parameters by 39.2% while maintaining low reconstruction error. Furthermore, the diverse size of the omic datasets may lead to some datasets dominating during training, diminishing the model’s generalizability and explainability. We develop a whole omic (view) dropout layer, which masks a complete omic layer based on a trainable hyperparameter. This provides a significant improvement in the model’s generalization, providing better reconstructions for cancer cell lines previously uncharacterized by specific omics (see ***Methods, Figure 1b)***. We then perform a multi-omic model explanation by calculating SHAP values ^15^ for all omic input features to assess their importance for the latent space integration and the reconstruction of omic features (see ***Methods***). This provides a systematic resource to explore potential nonlinear cancer genotype-phenotype associations.

Taken together, MOVE provides an unsupervised model that integrates all cancer cell line omics simultaneously. Using a 10-fold cross-validation strategy, MOVE’s reconstructed hold-out folds for CRISPR-Cas9 and drug responses were robustly correlated with the original data (mean Pearson’s r of 0.35 and 0.65, respectively) ***(Figure 2a, Supplementary Figure 3, Supplementary Table 5)***. MOVE performed better compared to a similar systematic supervised analysis designed to predict each CRISPR-Cas9 gene dependency either using core-omics (e.g., genomics, transcriptomics), only genomic, or only functionally related genes (mean best Pearson’s r 0.25) ^27^.

**Figure 2.**
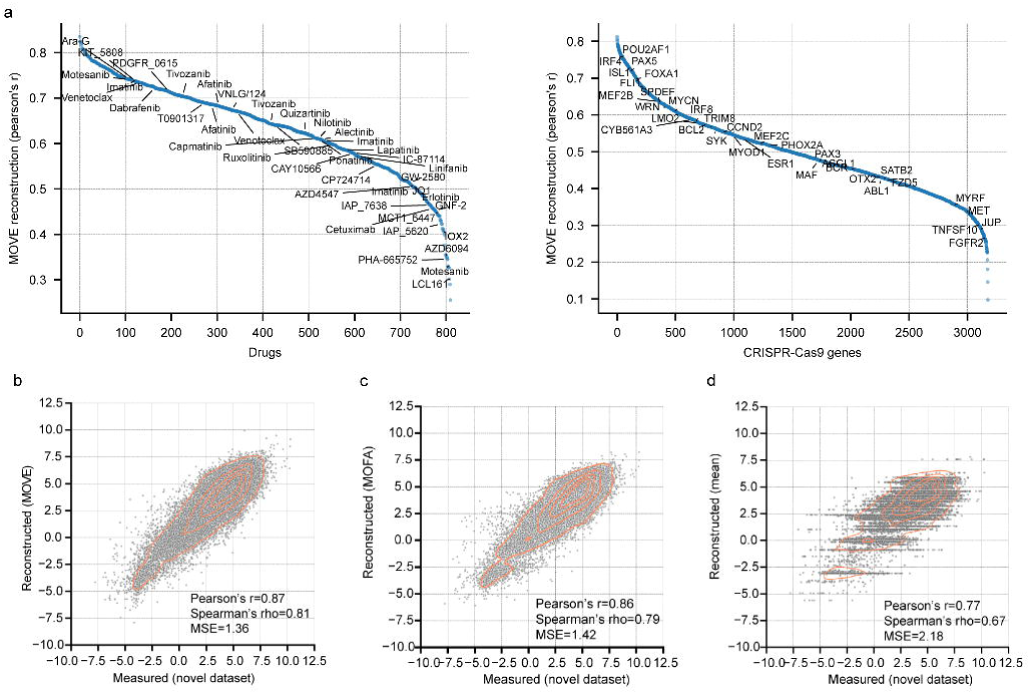
Reconstruction of MOVE for drug response and CRISPR-Cas9 datasets. **a)** MOVE reconstruction quality measured using a 10-fold cross-validation. After reconstructing all test folds, they are concatenated and the reconstruction quality score is calculated as the Pearson’s r between the reconstructed and measured values. Features ranked by their reconstruction quality are shown for the drug response (left) and the CRISPR-Cas9 (right) datasets. Duplicated drug names represent replicated screens for the same drug. Representative examples of strongly selective CRISPR-Cas9 and drug responses are labeled. **b)** MOVE’s partial dataset augmentation (missing value imputation) of drug IC50s compared to novel drug response screens. **c)** and **d),** similar to **b)**, using MOFA and mean imputed values, respectively.

### Evaluation of multi-omics synthetic data gene ration

A critical advantage of unsupervised deep-generative models is their ability to make robust synthetic data generation, i.e., reconstruct a dataset that is completely absent for specific cell lines (samples). This is particularly crucial given the pervasive gaps across the datasets, even in well-characterized models such as cancer cell lines ***(Figure 1a)***. Given that multi-omic profiling is both costly and labor-intensive, data-driven methodologies are key in prioritizing the most informative experiments. The robust validation of this is challenging as it requires independent, ideally large-scale datasets to validate the model’s predictions. We have delineated this over the following sections into specific benchmarks of increasing difficulty.

MOVE generates reconstructed versions of the input data matrices from the multi-omics latent space that was learned from the original data. Data reconstruction generates complete omic matrices, thus both handling missing values (partial dataset augmentation), and more importantly, reconstructing whole-omics for cell lines (full dataset augmentation) by using *in silico* generated omic measurements (at least two omics are required for a cell line to be considered in the model). For partial dataset augmentation, MOVE imputes incomplete features (e.g., measurements for certain proteins are sparse due to technical limitations) commonly found in mass-spectrometry-based proteomics data ^9,28^. A newly generated drug response dataset that is completely absent during model training was accurately reconstructed (IC50s, Pearson’s r 0.87, n=32,659) ***(Figure 2b)***, outperforming naïve mean imputation and unsupervised linear model ^8,12^ ***(Figure 2c and Figure 2d)***. Pronounced discrepancies between MOVE’s reconstruction and the original datasets revealed likely inaccurate experimental measurements. For example, the response to the MEK1/2 inhibitor trametinib was not consistent with replicate measurements and drugs with the same canonical target in the same cell line ***(Supplementary Figure 4a)***. Such discrepancies also spotlighted drugs (e.g., venetoclax) or classes of drugs (e.g., antiapoptotic inhibitors) for which no effective molecular biomarkers are available ***(Supplementary Figure 4b)***, underlining the challenge of devising reliable predictive models for their response. Additionally, proteomics is riddled with missing values ***(Supplementary Figure 5a)***, affecting more predominantly lowly abundant proteins. With the synthetic data, MOVE augmented the proteomics data by filling approximately 32% of the original matrix using information from all other omics, while preserving sample correlations with an independent proteomic dataset (CCLE ^29^) ***(Supplementary Figure 5b, Supplementary Table 6)***. Notably, MOVE effectively reconstructed the protein profiles of SMAD4 in cell lines characterized by SMAD4 gene deletions, which are typically associated with low SMAD4 gene expression and protein abundance ***(Supplementary Figure 5c)***. The MOVE-augmented proteomic matrix preserves the ability to identify protein interactions through protein pairwise correlations ^30^ ***(Supplementary Figure 5d)***. In contrast to the original matrix that has missing values, MOVE’s augmented protein matrix is complete and directly usable in downstream analysis, such as generalized linear models ^31^, which improved the recall of protein complex interactions ***(Supplementary Figure 5d)***.

For full dataset reconstruction, the synthetic proteomic data (n=78) generated by MOVE for cancer cell lines completely lacking proteomic measurements showed correlations with independent proteomic measurements similar to those of cell lines with proteomic measurements ***(Figure 3a)***. In terms of drug response, reconstructions for 107 overlapping drugs correlated robustly with measurements in an independent dataset (CTD2 ^32,33^) ***(Supplementary Figure 3b)***. Crucially, this shows the capacity of MOVE as a generative model for synthetic cancer cell line screening. We evaluated downstream analysis by comparing the original data matrices with the augmented ones. MOVE provided a 34.9% increase in the number of CRISPR-Cas9 cell line screens, and the augmented dataset provided an increase in statistical power to find genetic associations ***(Figure 3c, Supplementary Table 7)***. Gene essentiality specificity (Fisher’s skewness test) showed a moderate positive correlation (Pearson’s r = 0.52) between the synthetic CRISPR-Cas9 screened cell lines and the previously available screens ***(Figure 3d)***. Nonetheless, this correlation is likely underestimated due to the presence of potential outlier non-essential genes. MOVE accurately reconstructed gene dependencies, for example, BRAF dependency in BRAF gain-of-function mutant cancer cell lines ***(Figure 3e)***, and FLI1 dependency in cell lines harboring an EWSR1-FLI1 fusion gene ***(Figure 3f)***.

**Figure 3.**
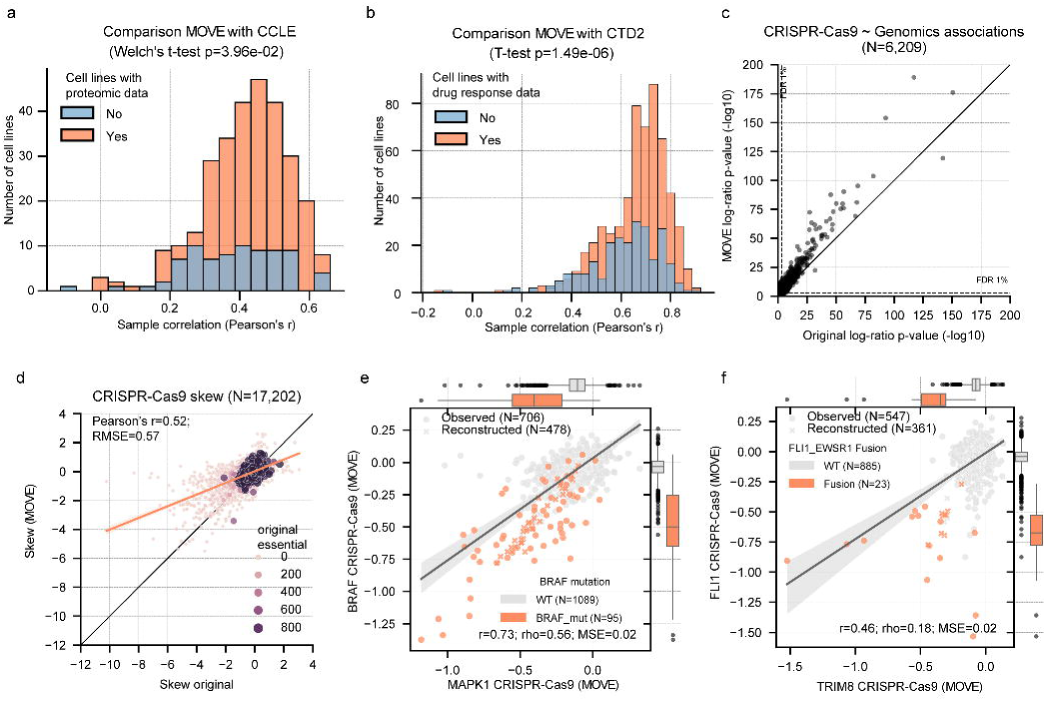
Multi-omics benchmark of MOVE. **a)** distribution of proteomics cancer cell lines correlation with an independent dataset (CCLE ^29^) grouped by whether the cancer cell line had proteomic data for the model training (orange) versus cell lines without any proteomics prior (light blue). **b)** distribution of cancer cell line correlations (Pearson’s r) between an independent drug response dataset (CTD2 ^32,33^) and the MOVE reconstructed dataset, grouped by whether the cancer cell line had prior availability of drug response in the datasets for the model training (orange) versus cell lines without drug response data (light blue). **c)** Log-ratio p-value of genetic associations with CRISPR-Cas9 gene essentiality with the original dataset (x-axis) and the augmented MOVE dataset (y-axis). **d)** Fisher skew test per gene across the original CRISPR-Cas9 dataset (x-axis) and the MOVE augmented dataset (y-axis). Dot size represents the number of cell lines that have the gene as essential (scaled log2 fold-change < -0.5) in the original dataset. **e)** correlation between BRAF and MAPK1 CRISPR-Cas9 gene essentialities using both previous measured (Observed) and newly synthetically reconstructed (Reconstructed). Gene essentiality scores are represented using copy-number corrected ^34^ log2 fold-changes scaled by the median of common essential (score = -1) and non-essential (score = 0) genes ^23^. Gene essentialities are also grouped according to the presence or absence of a BRAF mutation, mostly V600E gain-of-function mutations. **f)** CRISPR-Cas9 gene essentiality association with FLI1-EWSR1 fusion. **e)** and **f)** Box-and-whisker plots show 1.5 x interquartile ranges, centers indicate medians.

Taken together, these diverse examples demonstrate MOVE’s ability to perform both partial and full dataset augmentation across various datasets and laboratories. The generation of large-scale multi-omic datasets is both time and resource-intensive, thereby positioning MOVE as a valuable tool for *in silico* testing and prioritization of drug targets for experimental validation.

### Model interpretation reveals cancer cell states

To prioritize novel targets, a model needs to be explainable beyond producing reliable predictions. Hence, we used the SHAP ^15^ algorithm to calculate the feature importance, defined as the amount of contribution of each feature to the latent space (***Figure 1b,*** see ***Methods, Supplementary Table 8***). When grouping features by their corresponding omic datasets, we observed that metabolomics, drug response, and copy number variations exhibited the highest average feature importance (**Supplementary Figure 6a**). Regarding conditional features, although their average feature importance was modest, certain key features, such as TP53 mutation, growth rate, and tissue of hematopoietic and lymphoid origin emerged as highly significant, even when compared with other omic datasets ***(Figure 4a, Supplementary Table 8)***. This underscores the importance of incorporating conditional variables into the model. Features ranked in the top five from each omic dataset also validated the capacity of our approach to recover well-established molecular processes associated with cancer ***(Figure 4a)***, for example, CDKN2A copy number amplifications, as well as sensitivity to the SRC inhibitor, dasatinib. Interestingly, other less obvious features that were highly ranked shed light on previously less explored biological mechanisms. One specific example is the nicotinate and nicotinamide metabolism metabolite, 1-methylnicotinamide, that was calculated to be the most important feature in the metabolomics towards the multi-omics latent representation ***(Figure 4a)***. We observed a strong relation between increased 1-methylnicotinamide intracellular abundance and the overexpression of Nicotinamide N-Methyltransferase (NNMT) enzyme, which catalyzes the production of this metabolite ***(Supplementary Figure 6b)***. We also observed an association between 1-methylnicotinamide and the EMT state of cancer cell lines, as corroborated by the expression of VIM and CDH1 ^9^ ***(Supplementary Figure 6c and 6d)***. This confirms a recent single-cell study’s finding that the PC-9 non-small cell lung carcinoma line, which harbors an activating EGFR mutation, develops a cellular state resistant to EGFR inhibitors through expression of EMT markers as well as accumulation of 1-methylnicotinamide ^35^. While further experimental validation is necessary, this could pave the way for the identification of cancer cellular states underlying drug resistance.

**Figure 4.**
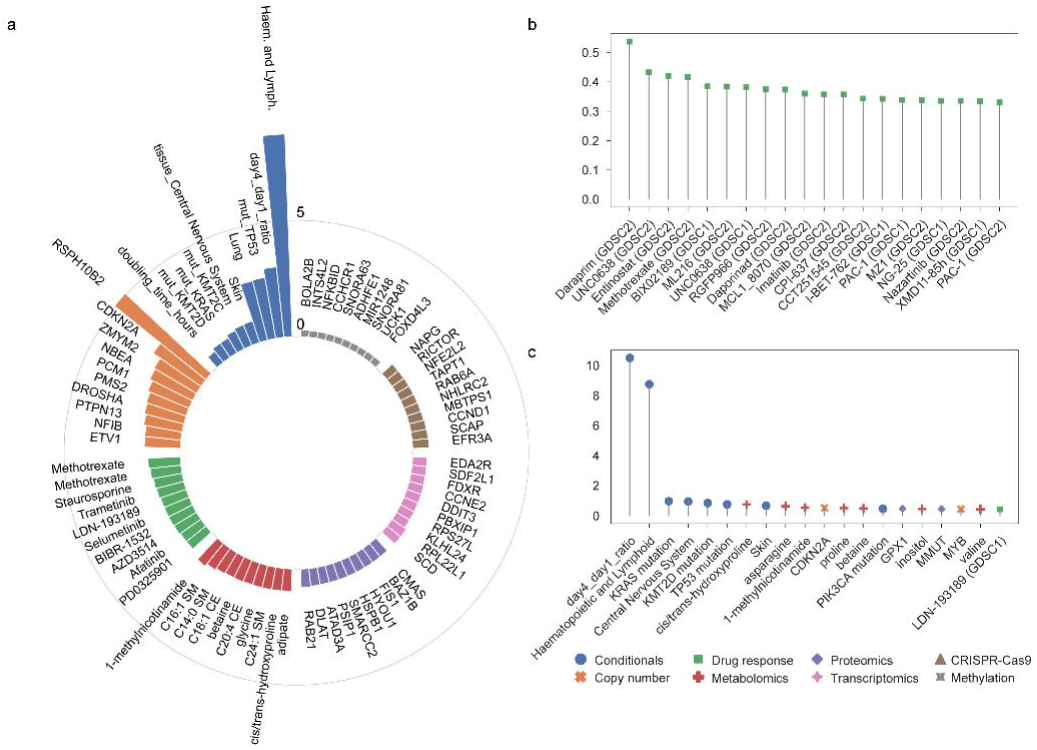
Model explanation of MOVE. **a)** top features from each omic layer that contribute the most to the multi-omic latent space. **b)** top drugs that have the highest feature importance from metabolite 1-methylnicotinamide. **c)** top features that contribute the most to the reconstruction of the drug response of Fulvestrant.

To delve deeper, we subsequently used the SHAP algorithm to calculate the feature importance specifically for the reconstruction of drug response, thereby facilitating the discovery of potential novel biomarkers (see ***Methods***). At the omic layer level, as expected, the drug response features themselves were the most important features on average ***(Supplementary Figure 7a, Supplementary Table 9)***. Notably, the conditionals emerged as the second most important omics, reflecting the critical role of tissue types, mutations, and growth rate in influencing drug responses ***(Supplementary Figure 7a)***. Centering on the metabolite 1-methylnicotinamide, drugs known to be EMT-related were ranked as the top drugs showing high feature importance from 1-methylnicotinamide ***(Figure 4b)***. All the top five drugs, except Daraprim which was not included in the dataset as an anti-cancer drug, were found to be related to EMT in recent studies. Specifically, UNC0638 ^36^, Entinostat ^37^ and BIX02189 ^38^ suppress EMT, while methotrexate ^39^ shows the ability to induce EMT. This finding suggests that the top-ranked drug Daraprim may also harbor a close relation to EMT, presenting a potential avenue for repurposing in cancer treatment. Other EMT-related features such as GPX1 protein intensity ^40^ also ranked as top features for Daraprim, indicating the potential to utilize other features in the list for the discovery of novel biomarkers for drug response ***(Figure 4c)***. Among the other top features for the top drugs, KRAS and KMT2D were consistently identified as being of high importance, and both of these genes have been implicated in EMT ^41,42^ ***(Supplementary Figure 7b-f)***.

Taken together, our findings suggest a broad association of 1-methylnicotinamide and EMT across hundreds of cancer cell lines with a potential role in drug resistance. While further assessment is needed to substantiate this, more generally, it unveils the possibility of using MOVE as a holistic model that integrates molecular and phenotypic data of cancer cells to investigate cancer cell states, drug resistance and their underlying mechanisms.

### Limitations and future work

The application of deep generative models, including MOVE, in cancer research is promising but comes with limitations, mainly that the restricted sample size impaired exploring more complex VAE designs. While the overall reconstruction of the datasets was robust, there are examples where it could be improved, particularly for proteomics where intrinsic data sparseness makes it more challenging for the model to train successfully. Thus, the addition of more characterized cancer models will likely allow to train better models and reduce reconstruction error. Future efforts should leverage multi-omic resources from cancer patients to enhance training by transfer learning or augmenting the sample pool if datasets can be integrated, and thereby potentially yield more generalizable predictions across different cancer cell models. SHAP analysis explains deep learning models, however there are still some obstacles in verifying the biological significance of certain highlighted features. These challenges could be associated with the inherent limitations of SHAP and Shapley values ^43^, thus additional research is required to ascertain the importance of these emphasized features.

## Conclusions

MOVE augmented the multi-omic profiles of 1,523 cancer cell lines by robustly filling in gaps in the existing experimental screens. Deep learning-based synthetic data generation can augment experimental screens by facilitating the creation of realistic datasets to guide experimental design and accelerate the validation of novel targets. Looking ahead, this model is readily adaptable to integrate other types of data modalities, such as imaging, further enabling the discovery of molecular/phenotype associations.

## Methods

### Cancer cell line multi-omic data collection

The aim was to assemble the most up-to-date and comprehensive molecular, phenotypic and cancer cell line sample information. All datasets were downloaded from the DepMap (https://depmap.org/) and the CellModelPassports (https://cellmodelpassports.sanger.ac.uk/) ^16^ portals, with the exception of the metabolomics data which were taken directly from the original publication supplementary materials ^20^. For reproducibility, all data used in this study are provided in a figshare repository.

We integrated genomics ^2,44^, transcriptomics ^19^, methylomics ^18^, proteomics ^9^, metabolomics ^20^, drug response ^18,21,22^, and CRISPR-Cas9 gene essentiality ^4,45^. This comprised a total of 1,994 cancer cell lines with at least two datasets available for each cell line. All datasets have been previously processed, normalized/scaled, and batch corrected in each of their individual publications addressing technical and design aspects important to each dataset (e.g., integration of CRISPR-Cas9 screens across different laboratories ^46^, driver mutations and copy number alterations, and gene expression samples from different datasets ^19^).

### Cancer cell line validation datasets

Three independent datasets were used in this study for validation, which were not used during the training phase. The first dataset presents the CCLE proteomic characterization of 375 cancer lines ^29^, of which 291 comprise the proteomic dataset ^9^ used for training. The second dataset represents novel drug response screens with the same platform as the drug screens used for training ^18,21,22^ that were obtained from the Genomics of Drug Sensitivity in Cancer (GDSC) portal (https://www.cancerrxgene.org/) ^47^ comprising a total of 32,659 IC50s measured across 313 unique drugs and 781 overlapping cancer cell lines. The third dataset is an independent drug response dataset (CTD2) ^32,33^, comprising a total of 545 drugs and 887 cancer cell lines, for which 106 and 575, respectively, overlap with the drug response data used for training ^18,21,22^.

### Data preprocessing

A total of seven datasets were considered: copy number (n=777 features); methylome (n=14,608); transcriptome (n=15,278); proteome (n=4,922); metabolome (n=225); drug response (n=810); and CRISPR-Cas9 gene essentiality (n=17,931). A total of 1,523 cancer cell lines were profiled with each cell line having at least two of these datasets.

For CRISPR-Cas9 gene essentiality, transcriptomic and methylomic feature reduction was performed to exclude lowly variable features. For gene essentiality, samples were scaled using essential and non-essential genes making their median per sample -1 and 0, respectively. Never essential genes were discarded, i.e., genes that do not have an essentiality profile lower than 50% of the median essential genes in at least one cell line were removed. For transcriptomics and methylomics, a standard deviation filter was applied. By taking the standard deviation of all genes across samples, a Gaussian mixture model (k=2) was fitted, identifying lowly variable genes and the rest. A standard deviation threshold was defined as the rightmost intercept of the two Gaussian distributions ***(Supplementary Figure 2a)***, and any gene with a standard deviation lower than that was discarded. Moreover, for the proteomic, drug response, metabolomic and CRISPR-Cas9 datasets, any feature with a missing rate higher than 85% was discarded. All datasets were standardized by z-score, except copy number. Missing values were replaced with 0 and their position in the original dataset was stored for use in the model (e.g., to exclude them from the loss functions). In addition to these seven datasets, specific gene mutations, fusion genes, microsatellite instability, growth rate, cancer and tissue type information were concatenated into a single matrix to be used as labels of the cancer cell lines.

### Multi-omics variational autoencoder (MOVE)

MOVE is a conditional multi-view variational autoencoder implemented using PyTorch (v2.0) ^48^. In the next section, we describe MOVE’s architecture, use of conditionals, dropout layer and SHAP explainability analysis.

#### Architecture

MOVE follows a traditional design of conditional VAEs ***(Figure 1b)***. For each of the seven datasets (views), an encoder is trained with multiple fully connected layers, which are all proportional to the number of input features of the dataset plus the number of labels (concatenated conditionals). First, a joint fully connected layer takes as input all the datasets concatenated and reduces them to a fixed number of joint latent dimensions. Different latent dimensions techniques were tested (e.g., product of experts), but concatenation obtained the smallest reconstruction loss. Then, the joint layer outputs two layers representing Gaussian distribution mean and variance. These are important for the regularization of the latent space and are used to sample the latent dimensions. Finally, the latent dimensions (z) are concatenated with the conditionals and provided to the encoders of each dataset. The latent dimensions have a similar but inverse architecture to the encoder. Highlights of MOVE are as follows.

#### Conditionals

We introduced a conditional architecture to enhance the model’s reconstruction performance and biological relevance. Conditionals (n=237) include key biological features, such as cancer driver mutations, tissue types, gene fusions, MSI status, and cell line growth rate. These were used in two moments: concatenated to each omic layer prior to encoding; and to the latent neurons before decoding. The conditional concatenation serves two crucial purposes: it contextualizes the input data within specific biological or genetic backgrounds, and it allows the decoder to generate condition-specific reconstructions of the data. The inclusion of conditionals offered several advantages. First, it ensured that the model was not merely capturing patterns within individual omic layers in isolation. Instead, complex interactions among multi-omic data and genomic and physiological variables were accounted for, facilitating a more holistic understanding of the underlying biological processes and phenomena. Second, by embedding these conditionals into the decoder, the model can generate data reconstructions contextualized to specific cell line conditions.

#### View dropout layer

A special dropout strategy, namely view dropout layer, was included in MOVE to both improve the model’s predictive power and interpretability. Unlike traditional dropout layers, which randomly set individual features to zero, the view dropout layer zeroes out all the input features pertaining to a single omic layer. This approach encouraged the model to reconstruct the data by learning the relationships among multiple omic layers, rather than relying on one specific omic layer. For example, in generating drug response predictions, the MOVE model could disproportionately emphasize the input drug response data, neglecting the potential contributions from other omic layers such as transcriptomic and proteomic data. By using the view dropout layer, we significantly improved the latent space cell line separation (***Figure 1c***, ***Supplementary Figure 1b, 8a-b***) and reconstruction for both the proteomic (***Figure 3a, Supplementary Figure 8c***) and drug response data (***Supplementary Figure 8d***). The dropout rate for this layer is controlled by the hyperparameter view_dropout, which was optimally set as 0.5 for the final model.

#### Model explanation via SHapley Additive exPlanations (SHAP)

For model explanation, we used the Python package SHAP^15^ (v0.42.1) with technical modifications to support the multi-omic data as the input to MOVE. Specifically, the GradientExplainer, which combines IntegratedGradient ^49^ and SmoothGrad ^50^, was used to calculate the changes of the gradients on the model’s output regarding its input to attribute an importance value to each feature. The SHAP calculation was performed in two ways.

First, SHAP was run to explain the encoder part of MOVE, treating the integrated latent dimensions as the output. The result contains SHAP values in a multidimensional array with shapes of (*N_latent_dim_, N_samples_, N_features_*), where each *N* represents the number of latent dimensions, samples and features, respectively. To achieve the global level feature importance for analysis, the multidimensional array was first taken as the absolute value to account for both positive and negative impact, and then summed across latent dimensions, followed by averaging by samples. This then resulted in a list of length *N_features_*, representing the overall feature importance contributing to the latent space (**Figure 4a**).

Second, SHAP was run to explain MOVE’s reconstruction of each omic dataset. Taking drug response as an example, similarly to explaining the latent space, the shape of the SHAP values is (*N_drug_, N_samples_, N_features_*), where *N_drug_* represents the number of drugs, and *N_samples_, N_features_* are described as above. In this analysis, the array was only averaged across samples, resulting in a 2D array of (*N_drug_, N_features_*), which measures the feature importance for each drug. The feature 1-methylnicotinamide metabolite was first selected and the drugs that had the highest SHAP values were analyzed to identify EMT-related drugs (***Figure 4b***). Other important features for the drugs of interest were then ranked by selecting the row of the drug and then ranking the features in the descending order (***Figure 4c***). Due to the limitation of the computational resource, 20% of the samples were randomly selected to compute the feature importance for reconstructing drug response and copy number datasets, while 20 samples were randomly selected for other omic datasets which have much larger number of dimensions.

Overall, the SHAP analysis allowed us to identify features that are important for the multi-omic latent dimension and for explaining the reconstruction of features, such as drug response. Feature importance aggregated across all the samples and output dimensions can be found in ***Supplementary Table 8*** and ***9***. More granular feature importance scores for each output dimension can be downloaded from the figshare repository provided in the *code and data availability* section.

#### Loss function

The loss function is the summation of three components: 1) Reconstruction error across all input datasets; 2) weighted variational Kullback-Leibler (KL) regularization term of the joint latent space ^51^; and 3) a contrastive loss using tissue types as labels:

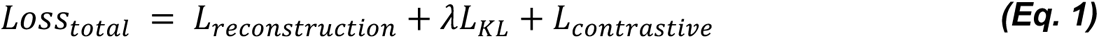

The reconstruction loss *L_reconstruction_* is defined as:

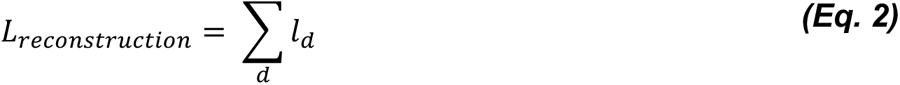

Where *l_d_* represents the reconstruction loss for dataset *d*, calculated using the mean squared error (MSE) ^52,53^.

In (***Eq.1)*** the *λ* is an optimized hyperparameter to weight for the KL divergence loss *L_KL_*, which calculates the KL divergence between the learned mean (*μ*) and variance (*σ*^2^) of the VAE and a standard normal prior distribution ^51^.

The last part of the loss function is a contrastive loss defined as:

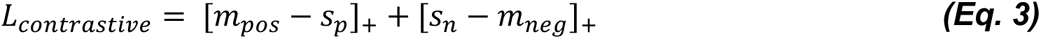

where *s_p_*and *s_n_* represents the cosine similarity between positive pairs and negative pairs, which are defined by whether two samples have the same tissue type. *m_pos_* and *m_neg_* are positive and negative margins, which are hyperparameters tuned as described in the section below.

#### Asymmetrical VAE

MOVE was also engineered with an asymmetrical structure to optimize model efficiency by reducing the number of parameters. Specifically, feature selection was conducted in a data-type-specific manner before the encoding process. For transcriptomic and methylation data, only features that exhibited high variability were selected as input to the model. Highly variable features were defined using a gaussian mixture model with two components fitted to the standard deviation of all features, thus capturing two distributions of lowly and highly variable features. The standard deviation threshold is defined as the biggest value at which the densities of the two distributions are equal, hence features with a standard deviation greater than 1.122 for transcriptomic data and 0.064 for methylation data are considered highly variable and selected as input for the encoder. For CRISPR-Cas9 data, gene knockouts that did not significantly impact any cell line, as indicated by a gene fitness score higher than -0.5 in every cell line, were excluded from the input layer. This targeted feature selection effectively reduced the model’s computational burden. Despite this reduction in input complexity, all available features were included during the decoding process to reconstruct the data. This asymmetrical design was chosen for its ability to maintain the model’s predictive and reconstructive capacities while streamlining its architecture.

#### Hyperparameters

The choice of hyperparameters ***(Table 1)*** was guided by an automatic optimization framework based on parallel trials (Optuna ^54^) and then manually adjusted ***(Supplementary Figure 2)***. For each run a stratified shuffle split is performed, stratifying by hematopoietic and lymphoid cell lines, leaving 20% of the samples for testing. A total of 600 trials were performed, where each trial was capped to 150 epochs.

**Table 1.** Optimized hyperparameters used for training MOVE.

| Parameter | Value |
| --- | --- |
| Number of epochs | 500 |
| Batch size | 256 |
| Learning rate | 3e-4 |
| Number of cross-validation folds | 3 |
| Feature missing rate threshold (%) | 0.85 |
| Latent dimensions of each view (%) | 0.25 |
| Number of joint latent dimensions | 200 |
| Hidden dimensions (%) | 0.7 |
| Dropout probability | 0.4 |
| View dropout probability | 0.3 |
| Weight of Kullback–Leibler (KL) loss term | 0.0001 |
| Optimizer | Adam |
| Weight decay | 5e-4 |
| Activation function | PReLU |
| Scheduler | Plateau |
| Scheduler threshold | 1e-4 |
| Scheduler factor | 0.6 |
| Scheduler patience | 7 |
| Scheduler minimum learning rate | 1e-7 |

### Linear factor analysis dimensionality reduction (MOFA)

For comparison, we systematically benchmarked our results with a linear multi-omics dimensionality reduction approach performed with MOFA ^8,12^. To make comparisons as close as possible and focus solely on methodological differences, the same data preprocessing was used for MOVE and MOFA. Similarly, the number of factors for MOFA was initially set to the same optimal number of joint latent dimensions, i.e., 200 ***(Table 1)***. However, this generated poorly performant results, e.g., poorly reconstructed views. Through manual exploration, the optimal number of factors was set to 100, which were automatically reduced during training to 97 by discarding factors with variance explained lower than 0.0001. Each view was scaled independently, and the model was run until it converged (convergence_mode=slow). The conditionals layer in MOVE contains both binary, e.g., mutations and tissue of origin, and continuous, e.g., cancer cell line growth rates and doubling times from independent studies, features. However, MOFA does not support a mixed distributed view, e.g., Gaussian and Bernoulli. Thus, we could not integrate the growth rate and doubling time in the conditional view, which apart from these two have only binary features and therefore the prior likelihood distribution was set to Bernoulli. These configurations produced the best multi-omic dimensionality reduction and view reconstruction using MOFA. The optimized model was saved as an HDF5 file and is also provided in the figshare repository.

### Protein-protein interaction co-abundance analysis

Protein-protein interactions (PPIs) were estimated using two methods to compare the ability of MOVE and the original proteomic matrix to recapitulate PPIs present in specific protein interaction resources datasets: CORUM ^55^, BioGRID ^56^ and STRING ^57^. The first method, Pearson’s R, has been previously used for this task ^30^. Due to its inherent limitations, e.g., its inability to account for confounding effects and data structure, a new method, similar to that described by Wainberg et al. ^31^, based on a generalized linear model (GLM) was developed. This method applies Cholesky’s Whitening transformation to proteomics data by using the inverse of its covariance matrix, which decorrelates samples and pushes data towards normality. This transformed data is then used in an ordinary least squares (OLS), whose calculated weights are a correlative metric between two proteins. Each method was calculated for every protein pair, with all pairs being ordered for each method by ascending p-value. Afterwards, a curve was drawn based on the cumulative sum of presence of that pair on a PPI set, either 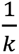 (presence) or 0 (absence), where *k* is the total number of present pairs. Thus, the better the method, the greater the AUC of the recall curve.

## Code and data availability

All code is available at https://github.com/QuantitativeBiology/PhenPred. All data were assembled from the Cancer DepMap and synthetic datasets generated are available for download at figshare:

● MOFA imputed datasets: https://doi.org/10.6084/m9.figshare.24420631;
● DepMap datasets: https://doi.org/10.6084/m9.figshare.24420580, https://doi.org/10.6084/m9.figshare.24420598;
● MOVE augmented datasets: https://doi.org/10.6084/m9.figshare.24562765;
● MOVE feature importance: https://doi.org/10.6084/m9.figshare.24473005;

## Supplementary Material

### Supplementary Figures

**Supplementary Figure 1.**
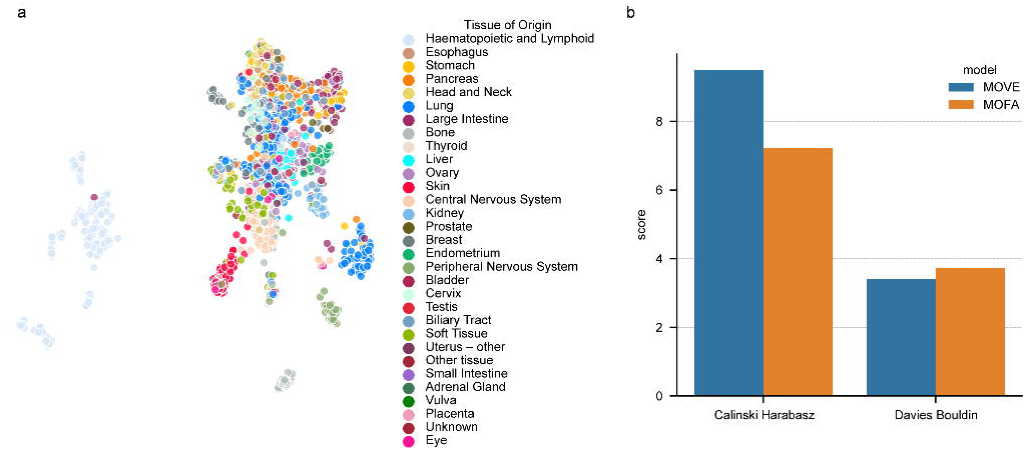
Latent space visualization comparison. **a)** UMAP representation of the trained MOFA joint latent space, where each dot represents a cancer cell line and is coloured according to its tissue of origin. **b)** comparison of cell line separations quantified by Calinski-Harabasz index (higher value indicates better) and Davies-Bouldin index (lower value indicates better).

**Supplementary Figure 2.**
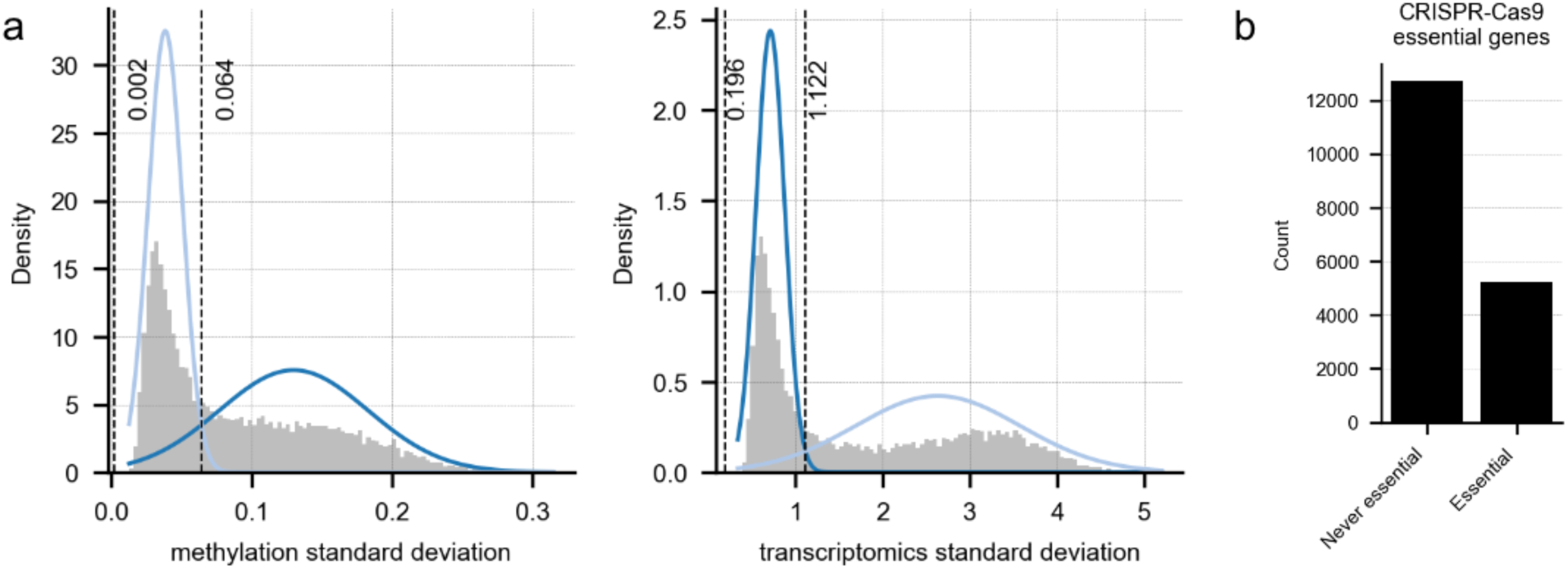
Feature filtering in MOVE. **a)** gaussian mixture model (k=2) trained to capture gene-promoter methylation (left panel) and gene expression (right panel) features that are lowly variable. Standard deviation thresholds at which the Gaussian mixtures have the same height, i.e., the probability of feature belonging to either distribution is the same, are represented by vertical lines. The rightmost threshold is used for feature selection for the encoders, specifically those that have a standard deviation smaller than the rightmost threshold are masked. **b)** CRISPR-Cas9 screens were scaled using previously reported essential and non-essential genes (see **Methods**), and genes with a scaled gene essentiality score lower than 50% of the median of essential genes (-1) were marked as 1, otherwise 0. Never essential genes denote those genes that were never found to have a scaled gene essentiality lower than the median of the essential genes, and therefore were filtered for the encoders.

**Supplementary Figure 3.**
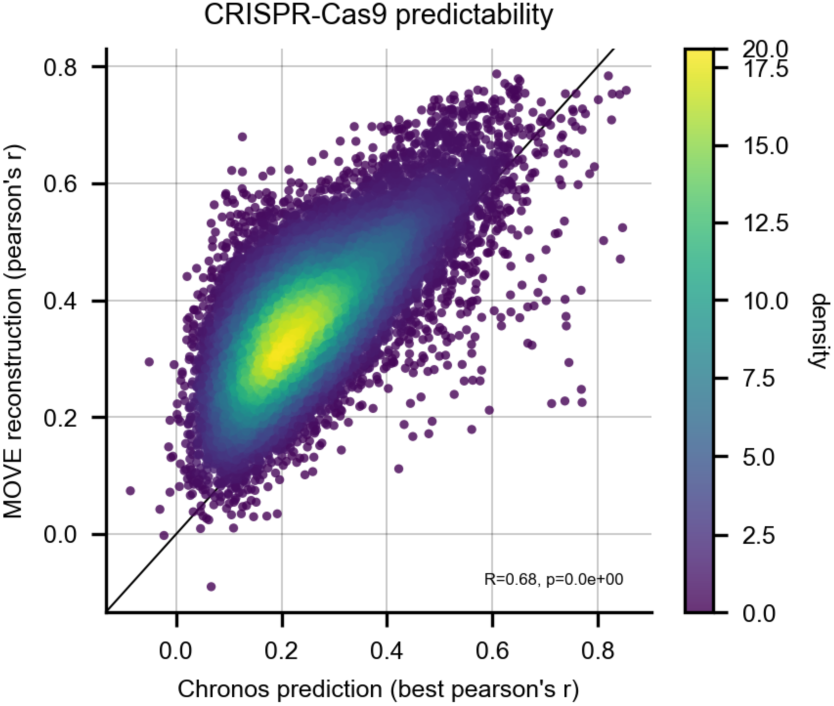
Comparison of MOVE’s reconstruction quality score against Chronos DepMap supervised prediction quality score calculated similarly (i.e., 10-fold cross-validation and Pearson’s correlation between reconstructed and measured) ^27^. For each gene, Chronos estimates three models (Core Omics, DNA-based and Related), and for this comparison the best Pearson’s r score per gene was selected.

**Supplementary Figure 4.**
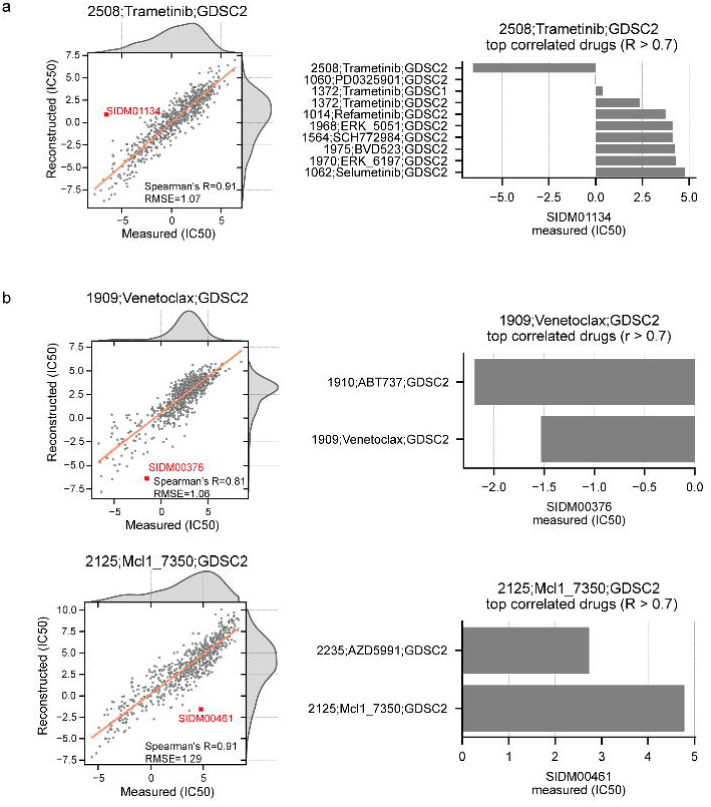
Drug response reconstruction benchmark. **a)** the biggest absolute difference between the reconstructed drug response data and the original dataset. In the left panel, a regression plot between the reconstructed drug response and the original measurements is shown, with the particular cell line, where the highest discrepancy was observed highlighted with a red cross. In the right panel, a barplot of the measured IC50 of the highlighted cell line is shown. The drugs were selected according to their IC50 correlation to the drug with the biggest discrepancy. **b)** similar to c), in this case, two representative drugs that target the antiapoptotic MCL1/BCL2 pathway were chosen and plotted among those with the highest discrepancies.

**Supplementary Figure 5.**
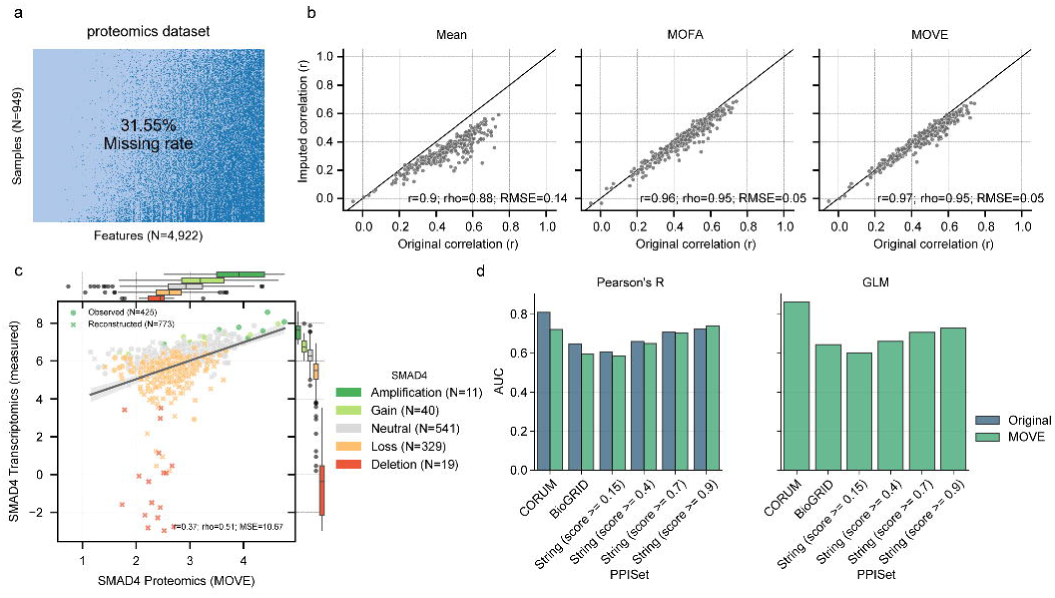
Proteomics missing value reconstruction. **a)** representation of missing values present in the proteomics dataset, where missing values are represented by dark-blue cells in the matrix, and observations with light-blue. **b)** comparison of the original (i.e., used for training MOVE) and imputed datasets with an independent proteomics dataset ^29^. Each dot represents a cancer cell line present in both the original and independent proteomics dataset and the correlation (Pearson’s r) between the two for every cell line is reported in the x-axis. In the y-axes the correlations are reported for the same cell lines, between the imputed matrices using the different methods and the independent dataset. **c)** correlation between measured SMAD4 gene expression and synthetic SMAD4 protein intensity by MOVE. Circle represents cell lines with original proteomic data (Observed) and cross represents cell lines with synthetic proteomic data (Reconstructed). Box-and-whisker plots show 1.5 x interquartile ranges, centers indicate medians. **d)** Bar plot of the area under the curve (AUC) of the recall curve when recapitulating protein-protein interactions (PPIs) present in a certain set. The PPI sets used are those from CORUM ^55^, BioGRID ^56^ and STRING ^57^ with 4 different score thresholds. The AUC was calculated using the p-value of either Pearson’s R correlation coefficient (left) or the effect sizes of the generalized linear model (right) ^31^. This last method cannot be computed on the original data since it has missing values.

**Supplementary Figure 6.**
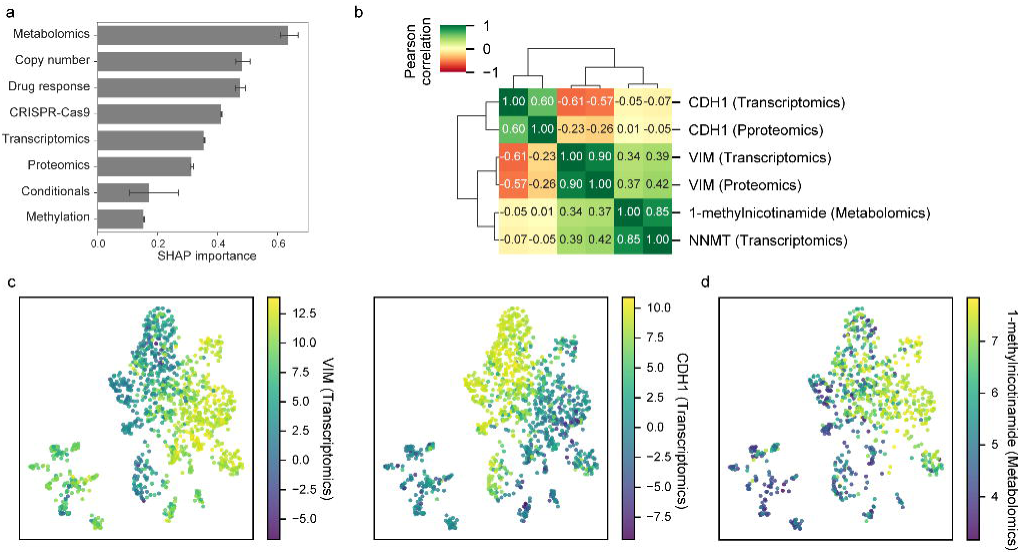
MOVE feature importance and joint latent space representations colored by specific markers, **a)** global feature importance grouped by omic datasets for constructing the joint latent space. **b)** correlation heatmap for some of the most predominant features of MOVE identified using the SHAP analysis. **c)** gene expression of canonical markers of epithelial mesenchymal transition (EMT), VIM (left) and CDH1 (right). **d)** similarly to b) but colored by the intracellular abundance of the 1-methylnicotinamide metabolite.

**Supplementary Figure 7.**
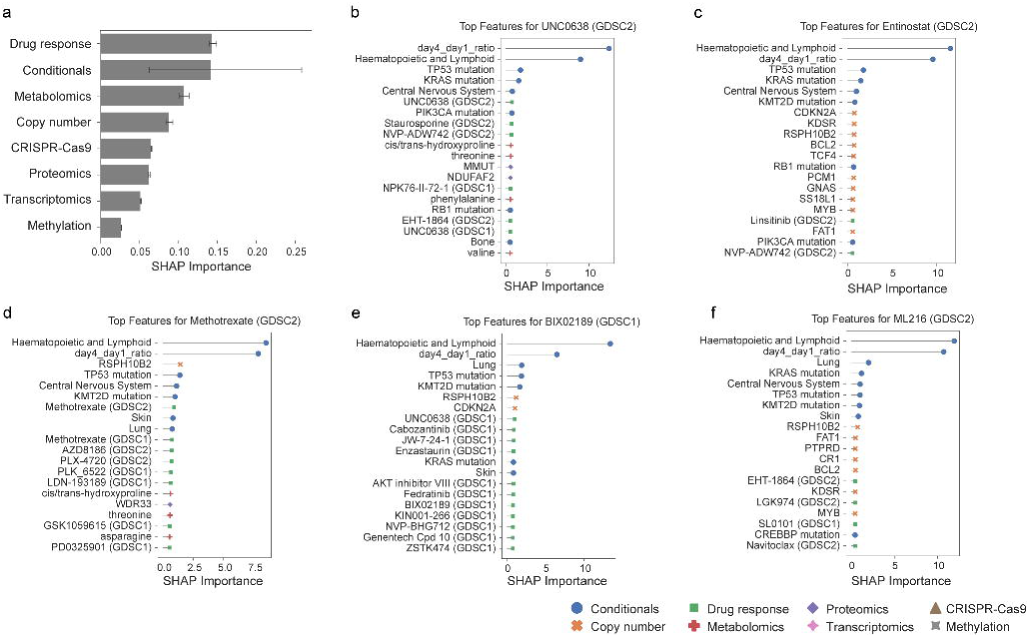
Feature importance calculated by SHAP values for the reconstruction of drug response. **a)** global feature importance grouped by omic datasets. **b-f**) top features for drugs that have the highest feature importance from 1-methylnicotinamide metabolite as shown in Figure 4b.

**Supplementary Figure 8.**
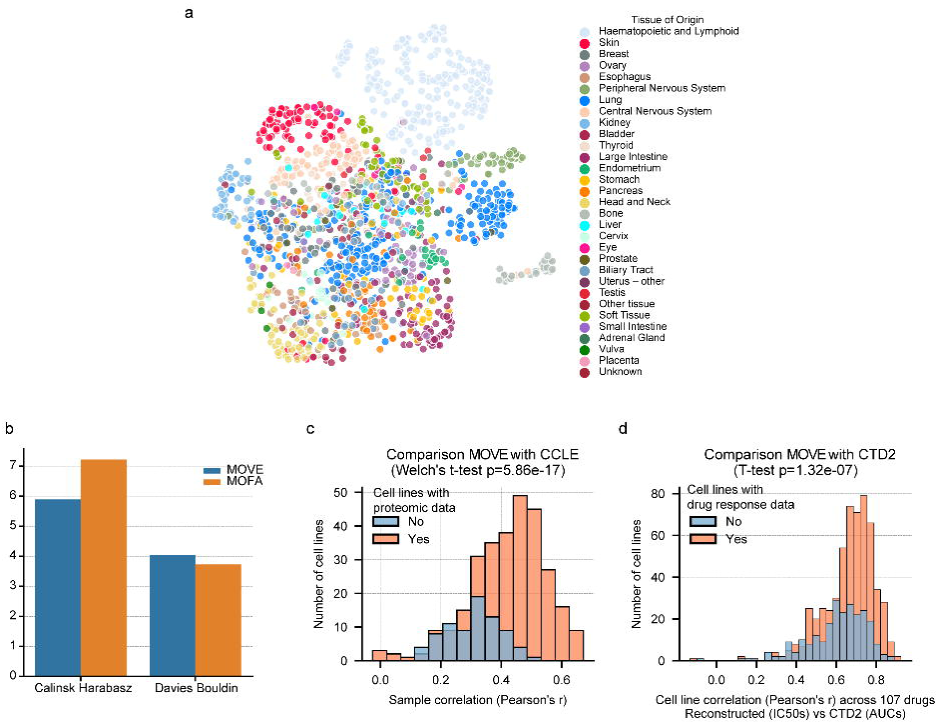
Evaluations of MOVE when view dropout is disabled. **a)** UMAP representation of the trained MOFA joint latent space. Similar to Figure 1c. **b)** comparison of cell line separations with MOFA. Similar to Supplementary Figure 1b. **c)** data reconstruction for proteomic data. Similar to Figure 3a. **d)** data reconstruction for drug response data. Similar to Figure 3b.

### Supplementary Tables

***Supplementary Table 1.*** Cancer cell line samplesheet and available datasets.

***Supplementary Table 2.*** Multi-omics MOVE latent dimensions.

***Supplementary Table 3.*** Conditional labels.

***Supplementary Table 4.*** Omic feature masks.

***Supplementary Table 5.*** Drug response and CRISPR-Cas9 10-fold cross-validation metrics.

***Supplementary Table 6.*** Sample-wise proteomics CCLE correlation using different imputation methods.

***Supplementary Table 7.*** CRISPR-Cas9 gene essentiality associations with genetic alterations.

***Supplementary Table 8.*** *Global level SHAP-based feature importance for explaining the latent space, aggregated across all the samples and latent dimensions*.

***Supplementary Table 9.*** *Global level SHAP-based feature importance for explaining the drug response, aggregated across all the samples and drugs*.

## Acknowledgments

We thank the Broad Institute and the Wellcome Sanger Institute for, through the Cancer Dependency Map consortium, making their data freely available and readily accessible to the scientific community and thereby enabling this work. This research was funded in part by the Wellcome Trust Grant 206194. For open access, the authors have applied a CC BY public copyright license to any Author Accepted Manuscript version arising from this submission.

## Author information

### Contributions

Z.C., S.A., A. R. B., M.D.S., C.P. and E.G implemented analyses. Z.C., S.A. and E.G. wrote the software. E.G. supervised and conceptualized the study. S.V., P.J.R., R.R.R., M.J.G, Q.Z. and E.G acquired funding and contributed to methodology. Z.C, Q.Z. and E.G wrote the manuscript. All authors have revised and approved the manuscript.

### Inclusion and ethics

All authors have committed to upholding the principles of research ethics and inclusion as advocated by the Nature Portfolio journals.

## References

1. Trastulla, L., Noorbakhsh, J., Vazquez, F., McFarland, J. & Iorio, F. Computational estimation of quality and clinical relevance of cancer cell lines. Mol. Syst. Biol. 18, e11017 (2022).

2. Garnett, M. J. et al. Systematic identification of genomic markers of drug sensitivity in cancer cells. Nature 483, 570–575 (2012).

3. Barretina, J. et al. The Cancer Cell Line Encyclopedia enables predictive modelling of anticancer drug sensitivity. Nature 483, 603–607 (2012).

4. Behan, F. M. et al. Prioritization of cancer therapeutic targets using CRISPR–Cas9 screens. Nature 568, 511–516 (2019).

5. Tsherniak, A. et al. Defining a Cancer Dependency Map. Cell 170, 564–576.e16 (2017).

6. Cai, Z., Poulos, R. C., Liu, J. & Zhong, Q. Machine learning for multi-omics data integration in cancer. iScience 25, 103798 (2022).

7. Argelaguet, R. et al. Multi-omics profiling of mouse gastrulation at single-cell resolution. Nature 576, 487–491 (2019).

8. Argelaguet, R. et al. MOFA+: a statistical framework for comprehensive integration of multi-modal single-cell data. Genome Biol. 21, 111 (2020).

9. Gonçalves, E. et al. Pan-cancer proteomic map of 949 human cell lines. Cancer Cell 40, 835–849.e8 (2022).

10. Eraslan, G., Simon, L. M., Mircea, M., Mueller, N. S. & Theis, F. J. Single-cell RNA-seq denoising using a deep count autoencoder. Nat. Commun. 10, 390 (2019).

11. Freeman, B. A. et al. MIRTH: Metabolite Imputation via Rank-Transformation and Harmonization. Genome Biol. 23, 184 (2022).

12. Argelaguet, R. et al. Multi-Omics Factor Analysis-a framework for unsupervised integration of multi-omics data sets. Mol. Syst. Biol. 14, e8124 (2018).

13. Boehm, J. S. et al. Cancer research needs a better map. Nature 589, 514–516 (2021).

14. Poulos, R. C., Cai, Z., Robinson, P. J., Reddel, R. R. & Zhong, Q. Opportunities for pharmacoproteomics in biomarker discovery. Proteomics 23, e2200031 (2023).

15. Lundberg, S. M. & Lee, S.-I. A Unified Approach to Interpreting Model Predictions. in Advances in Neural Information Processing Systems (eds. Guyon, I. et al.) vol. 30 (Curran Associates, Inc., 2017).

16. van der Meer, D. et al. Cell Model Passports-a hub for clinical, genetic and functional datasets of preclinical cancer models. Nucleic Acids Res. 47, D923–D929 (2019).

17. Dwane, L. et al. Project Score database: a resource for investigating cancer cell dependencies and prioritizing therapeutic targets. Nucleic Acids Res. 49, D1365–D1372 (2021).

18. Iorio, F. et al. A Landscape of Pharmacogenomic Interactions in Cancer. Cell 166, 740–754 (2016).

19. Garcia-Alonso, L. et al. Transcription Factor Activities Enhance Markers of Drug Sensitivity in Cancer. Cancer Res. 78, 769–780 (2018).

20. Li, H. et al. The landscape of cancer cell line metabolism. Nat. Med. 25, 850–860 (2019).

21. Picco, G. et al. Functional linkage of gene fusions to cancer cell fitness assessed by pharmacological and CRISPR-Cas9 screening. Nat. Commun. 10, 2198 (2019).

22. Gonçalves, E. et al. Drug mechanism-of-action discovery through the integration of pharmacological and CRISPR screens. Mol. Syst. Biol. 2020.01.14.905729 (2020) doi:10.1101/2020.01.14.905729.

23. Meyers, R. M. et al. Computational correction of copy number effect improves specificity of CRISPR-Cas9 essentiality screens in cancer cells. Nat. Genet. 49, 1779–1784 (2017).

24. Zampieri, G., Vijayakumar, S., Yaneske, E. & Angione, C. Machine and deep learning meet genome-scale metabolic modeling. PLoS Comput. Biol. 15, e1007084 (2019).

25. Lotfollahi, M., Wolf, F. A. & Theis, F. J. scGen predicts single-cell perturbation responses. Nat. Methods 16, 715–721 (2019).

26. Ashuach, T. et al. MultiVI: deep generative model for the integration of multimodal data. Nat. Methods 20, 1222–1231 (2023).

27. Dempster, J. M., Krill-Burger, J., Warren, A. & McFarland, J. Gene expression has more power for predicting in vitro cancer cell vulnerabilities than genomics. bioRxiv (2020) doi:10.1101/2020.02.21.959627.

28. Poulos, R. C. et al. Strategies to enable large-scale proteomics for reproducible research. Nat. Commun. 11, 3793 (2020).

29. Nusinow, D. P. et al. Quantitative Proteomics of the Cancer Cell Line Encyclopedia. Cell 180, 387–402.e16 (2020).

30. Gonçalves, E. et al. Widespread Post-transcriptional Attenuation of Genomic Copy-Number Variation in Cancer. Cell Syst 5, 386–398.e4 (2017).

31. Wainberg, M. et al. A genome-wide atlas of co-essential modules assigns function to uncharacterized genes. Nat. Genet. (2021) doi:10.1038/s41588-021-00840-z.

32. Seashore-Ludlow, B. et al. Harnessing Connectivity in a Large-Scale Small-Molecule Sensitivity Dataset. Cancer Discov. 5, 1210–1223 (2015).

33. Rees, M. G. et al. Correlating chemical sensitivity and basal gene expression reveals mechanism of action. Nat. Chem. Biol. 12, 109–116 (2016).

34. Iorio, F. et al. Unsupervised correction of gene-independent cell responses to CRISPR-Cas9 targeting. BMC Genomics 19, 604 (2018).

35. Oren, Y. et al. Cycling cancer persister cells arise from lineages with distinct programs. Nature 596, 576–582 (2021).

36. Liu, X.-R. et al. UNC0638, a G9a inhibitor, suppresses epithelial-mesenchymal transition-mediated cellular migration and invasion in triple negative breast cancer. Mol. Med. Rep. 17, 2239–2244 (2018).

37. Du, L., Xie, F., Han, H. & Zhang, L. Targeting SALL4 by Entinostat Inhibits the Malignant Phenotype of Gastric Cancer Cells by Reducing EMT Signaling. Anticancer Res. 43, 4389– 4401 (2023).

38. Park, S. J. et al. BIX02189 inhibits TGF-β1-induced lung cancer cell metastasis by directly targeting TGF-β type I receptor. Cancer Lett. 381, 314–322 (2016).

39. Ojima, T., Kawami, M., Yumoto, R. & Takano, M. Differential mechanisms underlying methotrexate-induced cell death and epithelial-mesenchymal transition in A549 cells. Toxicol. Res. 37, 293–300 (2021).

40. Meng, Q. et al. Abrogation of glutathione peroxidase-1 drives EMT and chemoresistance in pancreatic cancer by activating ROS-mediated Akt/GSK3β/Snail signaling. Oncogene 37, 5843–5857 (2018).

41. Pan, L.-N., Ma, Y.-F., Li, Z., Hu, J.-A. & Xu, Z.-H. KRAS G12V mutation upregulates PD-L1 expression via TGF-β/EMT signaling pathway in human non-small-cell lung cancer. Cell Biol. Int. 45, 795–803 (2021).

42. Zhang, Y. et al. Genome-wide CRISPR screen identifies PRC2 and KMT2D-COMPASS as regulators of distinct EMT trajectories that contribute differentially to metastasis. Nat. Cell Biol. 24, 554–564 (2022).

43. Marques-Silva, J. & Huang, X. Explainability is NOT a Game. arXiv [cs.AI] (2023).

44. Ghandi, M. et al. Next-generation characterization of the Cancer Cell Line Encyclopedia. Nature 569, 503–508 (2019).

45. Pacini, C. et al. Integrated cross-study datasets of genetic dependencies in cancer. Nat. Commun. 12, 1661 (2021).

46. Dempster, J. M. et al. Agreement between two large pan-cancer CRISPR-Cas9 gene dependency data sets. Nat. Commun. 10, 5817 (2019).

47. Yang, W. et al. Genomics of Drug Sensitivity in Cancer (GDSC): a resource for therapeutic biomarker discovery in cancer cells. Nucleic Acids Res. 41, D955–61 (2013).

48. Paszke, A. et al. Pytorch: An imperative style, high-performance deep learning library. Adv. Neural Inf. Process. Syst. 32, (2019).

49. Sundararajan, M., Taly, A. & Yan, Q. Axiomatic Attribution for Deep Networks. in Proceedings of the 34th International Conference on Machine Learning (eds. Precup, D. & Teh, Y. W.) vol. 70 3319–3328 (PMLR, 06--11 Aug 2017).

50. Smilkov, D., Thorat, N., Kim, B., Viégas, F. & Wattenberg, M. SmoothGrad: removing noise by adding noise. arXiv [cs.LG] (2017).

51. Asperti, A. & Trentin, M. Balancing Reconstruction Error and Kullback-Leibler Divergence in Variational Autoencoders. IEEE Access 8, 199440–199448 (2020).

52. Kingma, D. P. & Welling, M. Auto-Encoding Variational Bayes. arXiv [stat.ML] (2013).

53. Kingma, D. P. & Welling, M. An Introduction to Variational Autoencoders. arXiv [cs.LG] (2019).

54. Akiba, T., Sano, S., Yanase, T., Ohta, T. & Koyama, M. Optuna: A Next-generation Hyperparameter Optimization Framework. in Proceedings of the 25th ACM SIGKDD International Conference on Knowledge Discovery & Data Mining 2623–2631 (Association for Computing Machinery, 2019).

55. Ruepp, A., et al. CORUM: the comprehensive resource of mammalian protein complexes. Nucleic Acids Res. 36, D646–50 (2008).

56. Chatr-Aryamontri, A. et al. The BioGRID interaction database: 2015 update. Nucleic Acids Res. 43, D470–8 (2015).

57. Szklarczyk, D. et al. The STRING database in 2017: quality-controlled protein-protein association networks, made broadly accessible. Nucleic Acids Res. 45, D362–D368 (2017).

